# Innate Immune Signaling in Hearts and Buccal Mucosa Cells of Patients with Arrhythmogenic Cardiomyopathy

**DOI:** 10.1101/2023.07.25.550526

**Authors:** Carlos Bueno-Beti, Alessandro Tafuni, Stephen P. Chelko, Mary N. Sheppard, Ella Field, Jennifer Tollit, Imogen K Heenan, Annabelle Barnes, Matthew R. Taylor, Luisa Mestroni, Juan Pablo Kaski, Jeffrey E. Saffitz, Angeliki Asimaki

## Abstract

**Objectives:** We sought to determine if persistent innate immune signaling via NFκB occurs in cardiac myocytes in patients with arrhythmogenic cardiomyopathy and if this is associated with myocardial infiltration of pro-inflammatory cells expressing CCR2. We also determined if buccal mucosa cells from young subjects with inherited disease alleles exhibit NFκB signaling.

**Background:** NFκB signaling in cardiac myocytes causes disease in a mouse model of arrhythmogenic cardiomyopathy by mobilizing CCR2-expressing macrophages which promote myocardial injury and arrhythmias. Buccal mucosa cells exhibit pathologic features similar to those seen in cardiac myocytes in patients with arrhythmogenic cardiomyopathy.

**Methods:** We analyzed myocardium from arrhythmogenic cardiomyopathy patients who died suddenly or required cardiac transplantation. We also analyzed buccal mucosa cells from young subjects with inherited disease alleles. The presence of immunoreactive signal for RelA/p65 in nuclei of cardiac myocytes and buccal cells was used as a reliable indicator of active NFκB signaling. We also counted myocardial CCR2-expressing cells.

**Results:** NFκB signaling was seen in cardiac myocytes in 34 of 36 cases of arrhythmogenic cardiomyopathy but in none of 19 age-matched controls. Cells expressing CCR2 were increased in patient hearts in numbers directly correlated with the number of cardiac myocytes showing NFκB signaling. NFκB signaling also occurred in buccal cells in young subjects with active disease.

**Conclusions:** Patients with clinically active arrhythmogenic cardiomyopathy exhibit persistent innate immune responses in cardiac myocytes and buccal mucosa cells reflecting an inflammatory process that fails to resolve. Such individuals may benefit from anti-inflammatory therapy.

**CONDENSED ABSTRACT:** NFκB signaling in cardiac myocytes causes arrhythmias and myocardial injury in a mouse model of arrhythmogenic cardiomyopathy by mobilizing pro-inflammatory CCR2-expressing macrophages to the heart. Based on these new mechanistic insights, we analyzed hearts of arrhythmogenic cardiomyopathy patients who died suddenly or required cardiac transplantation. We observed active NFκB signaling in cardiac myocytes associated with marked infiltration of CCR2-expressing cells. We also observed NFκB signaling in buccal mucosa cells obtained from young subjects with active disease. Thus, anti-inflammatory therapy may be effective in arrhythmogenic cardiomyopathy. Screening buccal cells may be a reliable way to identify patients most likely to benefit.

**HIGHLIGHTS:**

- Inflammation likely contributes to the pathogenesis of arrhythmogenic cardiomyopathy but the responsible mechanisms and the roles of specific classes of immune cells remain undefined.
- NFκB signaling in cardiac myocytes is sufficient to cause disease in a mouse model of arrhythmogenic cardiomyopathy by mobilizing injurious myeloid cells expressing CCR2 to the heart.
- Here, we provide evidence of persistent NFκB signaling in cardiac myocytes and increased CCR2-expressing cells in hearts of patients with arrhythmogenic cardiomyopathy. We observed a close correlation between the number of cardiac myocytes with active NFκB signaling and the number of CCR2-expressing cells in patient hearts.
- We also provide evidence of active NFκB signaling in buccal mucosa cells associated with initial onset of disease and/or disease progression in young subjects with arrhythmogenic cardiomyopathy alleles.

## INTRODUCTION

Inflammation appears to contribute to the pathogenesis of arrhythmogenic cardiomyopathy (ACM), a familial non-ischemic heart muscle disease characterized by arrhythmias and progressive myocardial injury, but its precise role is undefined. Inflammatory infiltrates occur typically in the hearts of ACM patients and their abundance has been correlated with “hot phases” of the disease associated with more frequent arrhythmias and accelerated myocardial injury (1, 2), but whether inflammatory cells directly injure the heart or accumulate mainly in response to injury caused by other mechanisms is not known nor it is known if specific classes of immune cells play particular roles. We have previously reported that inhibition of nuclear factor κB (NFκB) signaling rescues the disease phenotype in a mouse model of ACM involving homozygous knock-in of a variant in the gene encoding the desmosomal protein, desmoglein-2 (*Dsg2*^*mut/mut*^ mice) (3). In subsequent studies, we observed that NFκB signaling in cardiac myocytes is sufficient to drive the disease phenotype in *Dsg2*^*mut/mut*^ mice (4). Using a genetic approach to selectively prevent activation of NFκB signaling in cardiac myocytes alone virtually eliminated myocardial degeneration and its replacement by fibrosis, preserved contractile function and greatly reduced arrhythmias (4). We also observed a 5-fold increase in the number of monocytes/macrophages expressing C-C motif chemokine receptor-2 (CCR2+ cells) in the hearts of *Dsg2*^*mut/mut*^ mice, but no increase after blocking NFκB signaling in cardiac myocytes (4). CCR2 is a G-protein coupled receptor for a monocyte chemo-attractant family that includes monocyte chemoattractant protein-1 (MCP-1, aka CCL2). CCR2+ macrophages have been implicated in adverse cardiac remodeling (5, 6) and fibrosis (7), and MCP-1 expression is increased in hearts of *Dsg2*^*mut/mut*^ mice and in ACM patient iPSC-cardiac myocytes (3, 8). Furthermore, genetic deletion of *Ccr2* greatly reduces myocardial injury and arrhythmias in *Dsg2*^*mut/mut*^ mice (4). Taken together, these results indicate that NFκB signaling in cardiac myocytes drives myocardial injury, contractile dysfunction and arrhythmias, at least in part, by mobilizing injurious pro-inflammatory CCR2+ macrophages to the heart.

In light of new mechanistic insights gained through studies of mouse models of ACM, we sought to determine if NFκB signaling is activated in cardiac myocytes in ACM patients and, if so, whether this is associated with accumulation of myocardial CCR2+ cells. Furthermore, based on previous studies showing that buccal mucosa cells from ACM patients exhibit characteristic changes in the distribution of desmosomal and gap junction proteins similar to those seen in ACM patient hearts (9, 10), we also sought to determine if NFκB signaling is activated in buccal mucosa cells obtained from young individuals with inherited ACM alleles. Evidence of active NFκB signaling was indicated by the presence of immunoreactive signal for RelA/p65, the heterodimeric binding partner of NFκB, in nuclei of cardiac myocytes or buccal mucosa cells, whereas cytoplasmic signal alone meant NFκB was not activated. Using this approach, we observed active NFκB signaling in cardiac myocytes in hearts from ACM patients associated with a marked increase in CCR2+ cells. There was a close correlation between the number of myocardial nuclei showing positive RelA/p65 signal and the number of myocardial CCR2+ cells in each case. We also observed NFκB signaling in buccal mucosa cells from young ACM gene carriers which correlated with disease activity and/or the first onset of clinical manifestations of disease.

## METHODS

### Ethical approval

This study was approved by the UK National Health Service Research Ethics Committee (hearts: 17/LO/0747, buccal smears: 17/LO/0840). Informed consent was provided by next-of-kin at the time of autopsy and by parents/guardians of children whose buccal cells were sampled.

### Myocardial samples

Myocardial samples were analyzed from 36 ACM patients and 19 control subjects. Clinical, diagnostic and genetic information is shown in Table 1. Samples were obtained at autopsy from 29 ACM patients who had died suddenly, from explanted hearts of 6 ACM patients at heart transplantation, and from an endomyocardial biopsy performed in one ACM patient during internal defibrillator placement. Control myocardial samples came from age-matched subjects who died from non-cardiac causes and had no cardiovascular disease at autopsy (Table 1).

**Table 1:** Clinical, genetic and immunohistochemical data of myocardial samples.

| **Study No** | **Clinical/post-mortem data** | **Age at death** | **Sex** | **Genetic data** | **RelA/p65 signal location** |
| --- | --- | --- | --- | --- | --- |
| **Controls** | **Cause of death** |  |  |  |  |
| CTR1 | Homicide | 37 | M | NA | Cytoplasmic |
| CTR2 | Drowning | 29 | M | NA | Cytoplasmic |
| CTR3 | Electrocution | 24 | M | NA | Cytoplasmic |
| CTR4 | Drug overdose | 20 | F | NA | Cytoplasmic |
| CTR5 | Neck compression | 40 | M | NA | Cytoplasmic |
| CTR6 | Homicide | 34 | M | NA | Cytoplasmic |
| CTR7 | Drug overdose | 30 | F | NA | Cytoplasmic |
| CTR8 | Suicide | 21 | M | NA | Cytoplasmic |
| CTR9 | Drug overdose | 21 | M | NA | Cytoplasmic |
| CTR10 | Drug overdose | 21 | M | NA | Cytoplasmic |
| CTR11 | Drug overdose | 35 | M | NA | Cytoplasmic |
| CTR12 | Drug overdose | 28 | M | NA | Cytoplasmic |
| CTR13 | Drug overdose | 26 | M | NA | Cytoplasmic |
| CTR14 | Suicide | 30 | M | NA | Cytoplasmic |
| CTR15 | Head injury | 32 | M | NA | Cytoplasmic |
| CTR16 | Drowning | 19 | M | NA | Cytoplasmic |
| CTR17 | Motor vehicle accident | 49 | M | NA | Cytoplasmic |
| CTR18 | Pulmonary embolism | 29 | M | NA | Cytoplasmic |
| CTR19 | Motor vehicle accident | 23 | F | NA | Cytoplasmic |
| **ACM** |  |  |  |  |  |
| ACM1 | SCD; PM Dx: RV-dom ACM | 37 | M | *PKP2*; c.2198_2202 delACACC | Nuclear |
| ACM2 | SCD; PM Dx: RV-dom ACM | 21 | M | *PKP2*; Gln133X | Nuclear |
| ACM3 | SCD; PM Dx: RV-dom ACM | 20 | M | *PKP2*; c.253_256delGAGT | Nuclear |
| ACM4 | SCD; PM Dx: RV-dom ACM | 34 | M | *PKP2*; Ser688Pro | Nuclear |
| ACM5 | SCD; PM Dx: biventricular ACM | 20 | M | *PKP2*; 1216delG | Cytoplasmic |
| ACM6 | SCD; PM Dx: biventricular ACM | 21 | M | *PKP2*; 2013delC | Nuclear |
| ACM7 | SCD; PM Dx: biventricular ACM | 26 | F | *DSP*; c.4395T>G | Nuclear |
| ACM8 | SCD; PM Dx: biventricular ACM | 28 | M | *DSP*; c.5269C>T | Nuclear |
| ACM9 | biopsy at ICD implantation; Dx: biventricular ACM | Currently 35 | F | *DSP*; c.3045del | cytoplasmic |
| ACM10 | SCD; PM Dx: LV-dom ACM | 36 | F | *DSP*; R1113X | Nuclear |
| ACM11 | SCD; PM Dx: LV-dom ACM | 28 | F | *DSP*; c.5318del | Nuclear |
| ACM12 | SCD; PM Dx: biventricular ACM | 29 | M | *DSG2*; 2036delG | Nuclear |
| ACM13 | SCD; PM Dx: biventricular ACM | 18 | M | *DSC2*; G371fsX378 | Nuclear |
| ACM14 | SCD; PM Dx: biventricular ACM | 34 | M | *DSC2*; E896fsX900 | Nuclear |
| ACM15 | SCD; antemortem Dx: Naxos disease | 21 | M | *JUP*; 2157del2, homozygous | Nuclear |
| ACM16 | SCD; PM Dx: biventricular ACM | 21 | M | *PKP2*; R735Q & *DSP*; R270X | Nuclear |
| ACM17 | SCD; PM Dx: biventricular ACM | 32 | M | *DSG2*; M1I & *DSG2*; I333T | Nuclear |
| ACM18 | SCD; PM Dx: biventricular ACM | 20 | M | *TMEM43*; c.1073C>T | Nuclear |
| ACM19 | SCD; PM Dx: biventricular ACM | 45 | M | *TMEM43*; c.1073C>T | Nuclear |
| ACM20 | SCD; PM Dx: RV-dom ACM | 20 | M | *FLNC*; c.7552-1 G>A | Nuclear |
| ACM21 | SCD; PM Dx: RV-dom ACM | 36 | M | *FLNC*; c.6779A>G | Nuclear |
| ACM22 | transplant; Dx: biventricular ACM + HF | NA | NA | *PLN*; R14del | Nuclear |
| ACM23 | transplant; Dx: biventricular<br>ACM + HF | NA | NA | *PLN*; R14del | Nuclear |
| ACM24 | transplant; Dx: biventricular<br>ACM + HF | NA | NA | *PLN*; R14del | Nuclear |
| ACM25 | transplant; Dx: biventricular<br>ACM + HF | NA | NA | *PLN*; R14del | Nuclear |
| ACM26 | transplant; Dx: biventricular<br>ACM + HF | NA | NA | *PLN*; R14del | Nuclear |
| ACM27 | SCD; antemortem Dx: ACM | 37 | M | NA | Nuclear |
| ACM28 | SCD; PM Dx: biventricular ACM | 23 | F | NA | Nuclear |
| ACM29 | SCD; PM Dx: biventricular ACM | 37 | M | NA | Nuclear |
| ACM30 | SCD; PM Dx: RV-dom ACM | 41 | F | NA | Nuclear |
| ACM31 | SCD; PM Dx: RV-dom ACM | 20 | M | NA | Nuclear |
| ACM32 | SCD; antemortem Dx: ACM | 40 | M | NA | Nuclear |
| ACM33 | SCD; PM Dx: biventricular ACM | 24 | M | NA | Nuclear |
| ACM34 | SCD; PM Dx: biventricular ACM | 43 | M | NA | Nuclear |
| ACM35 | SCD; PM Dx: biventricular ACM | 33 | M | NA | Nuclear |
| ACM36 | transplant; Dx: biventricular<br>ACM + HF | Currently<br>47 | M | No pathogenic variants<br>detected on panel testing | Nuclear |

### Buccal mucosa samples

Buccal mucosa cells were obtained from 28 young individuals (ages 5-17 years) using a soft cytological brush as previously reported (9, 10). Clinical, diagnostic and genetic information is shown in Table 2. These subjects were either ACM probands or had a family history of ACM (21 were carriers of ACM disease alleles) and were being followed clinically at Great Ormond Street Hospital. Buccal cells were obtained from: 12 pre-clinical carriers of variants who had never shown clinical disease; 3 at the clinic visit following the first clinical manifestation of disease; 9 with a quiescent clinical picture at the time cells were obtained (termed “Stable” in Table 2); and 4 sampled during or shortly after clinical exacerbation of disease (“Hot Phase” in Table 2). Buccal smears from 22 children (ages 1-17 years) without clinical signs or family history of cardiomyopathy were used as negative controls.

**Table 2:** Clinical, Genetic and Immunocytochemical Data for Buccal Smear Samples.

| Study No | Age | Sex | Genetic data | Clinical status at time of sampling | Disease status | RelA/p65 signal distribution |
| --- | --- | --- | --- | --- | --- | --- |
| pACM1 | 13 | F | *PKP2*; c.148_151delACAG | Normal cardiac investigations to date | Preclinical | Cytoplasmic |
| pACM2 | 12 | F | *DSP*; c.7756C>T | Normal cardiac investigations to date | Preclinical | Cytoplasmic |
| pACM3 | 11 | M | *DSP*; Arg84X | Borderline left axis deviation; normal cardiac investigations | Preclinical | Cytoplasmic |
| pACM4 | 16 | M | *PKP2*; Thr50Ser*61 | Normal cardiac investigations to date | Preclinical | Cytoplasmic |
| pACM5 | 16 | F | *PKP2*; Thr50Ser*61 | Normal cardiac investigations to date | Preclinical | Cytoplasmic |
| pACM6 | 18 | F | *PKP2*; Thr50Ser*61 | Normal cardiac investigations to date | Preclinical | Cytoplasmic |
| pACM7 | 11 | M | *DSP*; c.1755_1756insA | Normal cardiac investigations to date | Preclinical | Cytoplasmic |
| pACM8 | 8 | M | *DSP*; c.1755_1756insA | Normal cardiac investigations to date | Preclinical | Cytoplasmic |
| pACM9 | 15 | M | *PKP2*; c.337-2A>T | Normal cardiac investigations to date | Preclinical | Cytoplasmic |
| pACM10 | 12 | F | *DSP*; Trp180Ter | Sibling of pACM22; small voltages in ECG leads; normal cardiac investigations | Preclinical | Cytoplasmic |
| pACM11 | 15 | F | *DSP*; p.Arg2160Ter | Normal cardiac investigations to date | Preclinical | Cytoplasmic |
| pACM12 | 5 | F | *DSP*; Gln 2692* homozygous | Normal cardiac investigations to date; extracardiac features include palmoplantar keratoderma, nail dystrophy, sparse hair and keratosis follicularis | Preclinical | Cytoplasmic |
| pACM13 | 16 | F | *DSP*; Arg2160X | Intraventricular conduction delay in V4 and early repolarization inferiorly and in V4-V6 compared to previous clinical investigations | Initial disease onset | Nuclear |
| pACM14 | 17 | M | *DSP*; Arg2160X | Early repolarization inferiorly and in V5, V6; SAEKG showed late potentials in one vector; moderate RV dilatation on cMRI compared to previous investigations | Initial disease onset | Nuclear |
| pACM15 | 17 | F | no pathogenic variants detected on gene panel | Evaluated after developing exertional syncope and light headedness associated with hyperventilation; T-wave inversions in V1-V3; positive SAEKG in 2 vectors; frequent ectopy | Initial disease onset | Nuclear |
| pACM16 | 12 | F | *DSP*; Leu622Pro | Previous diagnosis of ACM and erythrokeratoderma; frequent ectopy; biventricular dilatation; mildly impaired systolic function | Stable | Cytoplasmic |
| pACM17 | 14 | M | Not tested | Previous diagnosis of ACM; NSVT episodes on Holter monitoring. | Stable | Cytoplasmic |
| pACM18 | 15 | M | No pathogenic variants detected on gene panel | Previous diagnosis of ACM; ventricular ectopy on exertion; focal LGE on cMRI involving inferoseptal LV on cMRI; symptomatic narrow complex tachycardia on loop recorder | Stable | Cytoplasmic |
| pACM19 | 17 | M | No pathogenic variants detected on gene panel | Previous diagnosis of ACM; ventricular ectopy; exertional VT; borderline LV dilatation; impaired systolic function | Stable | Cytoplasmic |
| pACM20 | 17 | M | No pathogenic variants detected on gene panel | Previous diagnosis of ACM; NSVT; abnormal repolarization on resting ECG | Stable | Cytoplasmic |
| pACM21 | 17 | F | No pathogenic variants detected on gene panel | Previous diagnosis of ACM; positive SAEKG in 3 vectors; SVT on Holter monitoring; LGE in mid-wall basal anterior septum on cMRI | Stable | Cytoplasmic |
| pACM22 | 17 | M | *DSP*; Trp180Ter | Sibling of pACM10; previous diagnosis of ACM; right axis deviation; subendocardial LGE on cMRI involving LV free wall; normal LV volume with preserved systolic function | Stable | Cytoplasmic |
| pACM23 | 11 | M | *DSP*; c.7576delAAGA and *DSP*; c.6577G>A | Previous diagnosis of ACM; frequent ventricular ectopy; NSVT; previous sustained VT and mildly impaired LV and RV global systolic function | Stable | Cytoplasmic |
| pACM24 | 16 | M | *PKP2*; Thr50Ser*61 | Previous diagnosis of ACM; RV internal dimensions at upper limit of normal; SAEKG positive in 2 vectors | Stable | Cytoplasmic |
| pACM25 | 10 | F | *PKP2*; Lys859Asn, homozygous | Severe biventricular dilatation; impaired systolic function; frequent ectopy; VT; ICD in situ; on active cardiac transplant list | Hot phase | Nuclear |
| pACM26 | 17 | F | Results pending | Previous diagnosis of ACM with multifocal arrhythmia ablation in 2016; at time of buccal sampling, patient exhibited palpitations, dizziness and atrial tachycardia | Hot phase | Nuclear |
| pACM27 | 9 | F | *PKP2*; c.2489+1 G>A | Previous diagnosis of ACM; at time of buccal sampling, patient was being seen in clinic after a recent episode of palpitations, pallor and perioral cyanosis; ventricular triplets on ambulatory ECG monitoring | Hot phase | Nuclear |
| pACM28 | 8 | F | *SCN5A*; Arg222Gln | Previous diagnosis of ACM with LV dilatation and impaired global systolic function; at time of buccal sampling, patient showed >1000 PVCs/hour on ambulatory ECG monitor prompting ICD implantation | Hot phase | Nuclear |

### Immunoperoxidase staining of myocardial samples

Formalin-fixed, paraffin-embedded left ventricular myocardium from all ACM patients and controls was analyzed by immunoperoxidase staining using a primary antibody against RelA/p65. Sections (5μm thick) were deparaffinized, dehydrated, rehydrated and exposed to 3% hydrogen peroxide solution for 10 min to block endogenous peroxidase activity. Sections were then incubated first with blocking solution for 1 hr and then with a rabbit polyclonal anti-RelA antibody (LS-B653; LSBiosciences) overnight at 4°C. The following day, sections were incubated with donkey secondary antibody conjugated to horseradish peroxidase enzyme (1 hr). Peroxidase-conjugated antibodies were detected by the 3,3’-diaminobenzidine (DAB) substrate kit. Heart tissue was counterstained with Mayer’s hematoxylin. Bright field images were taken with a Nikon eclipse 80i microscope and recorded with Nikon DS-Fi1 camera.

### Immunofluorescence staining for CCR2

When sufficient tissue was available, additional myocardial sections were analyzed by immunofluorescence staining using a primary antibody against CCR2. Sections (5 μm thick) were deparaffinized, dehydrated, rehydrated and boiled in citrate buffer (pH=6.0) for 11 min. They were then first incubated with blocking solution for 1 hr and then with a mouse monoclonal anti-human CCR2 antibody (MAB150-SP; R&D Systems) overnight at 4°C. The following day, sections were incubated with anti-mouse IgG Cy3-labeled secondary antibody and mounted with ProLong Gold. Images were recorded using a Nikon A1R confocal microscope. The number of cells showing strong immunofluorescent signal was counted in 5 randomly selected fields and expressed as number of cells per mm^2^ section area.

### Immunocytochemistry of buccal cells

Buccal cells were smeared on glass slides and fixed by M-FIX spray (Merck Millipore). They were then incubated first with blocking solution for 1 hr and then with a rabbit polyclonal anti-RelA antibody (8242; Cell Signaling Technology) overnight at 4°C. The following day, samples were incubated with anti-rabbit IgG Cy3-labelled secondary antibody, counterstained with 4′,6-diamidino-2-phenylindole (DAPI) and mounted with ProLong Gold. Images were recorded using a Nikon A1R confocal microscope.

### Statistical analysis

Prism software (Version 9.2.0 (283); GraphPad Software Inc., San Diego, CA, USA) was used for the statistical analysis of the CCR2+ cell data. The normal distribution of the data was assessed by Kolmogorov–Smirnov test. Normally distributed data was analyzed for differences by 1-way analysis of variance (ANOVA) and the Newman–Keuls post-test for multiple comparisons. Pearson’s correlation coefficient (r) was used to assess the linear correlation between CCR2+ cells and RelA+ myocyte nuclei. Data are presented as the mean ± the standard error of the mean (SEM). A p-value <0.05 was deemed significant.

### Data availability

Additional data will be provided by the corresponding author upon request.

## RESULTS

### NFκB signaling is activated in cardiac myocytes in ACM patients

Strong immunoperoxidase signal for RelA/p65 was seen in cardiac myocyte nuclei in 34 of 36 ACM patient samples, but in none of the 19 control samples. Representative examples are shown in **Figure 1A**. Nuclear signal indicating activation of NFκB occurred in ACM patients with variants in all 5 desmosomal genes as well as variants in *FLNC, TMEM43* and *PLN* (Table 1). The two ACM cases with no RelA/p65 nuclear signal (ACM5 and ACM9 in Table 1) had variants in *PKP2* and *DSP*, respectively. It is not clear why these cases showed no evidence of activation of NFκB in cardiac myocytes. Nine ACM cases came from individuals who died suddenly out-of-hospital in whom the diagnosis of ACM was made on the basis of autopsy findings. No genetic screening information was available for these cases but they all showed clear evidence of active NFκB signaling in cardiac myocytes.

**Figure 1:**
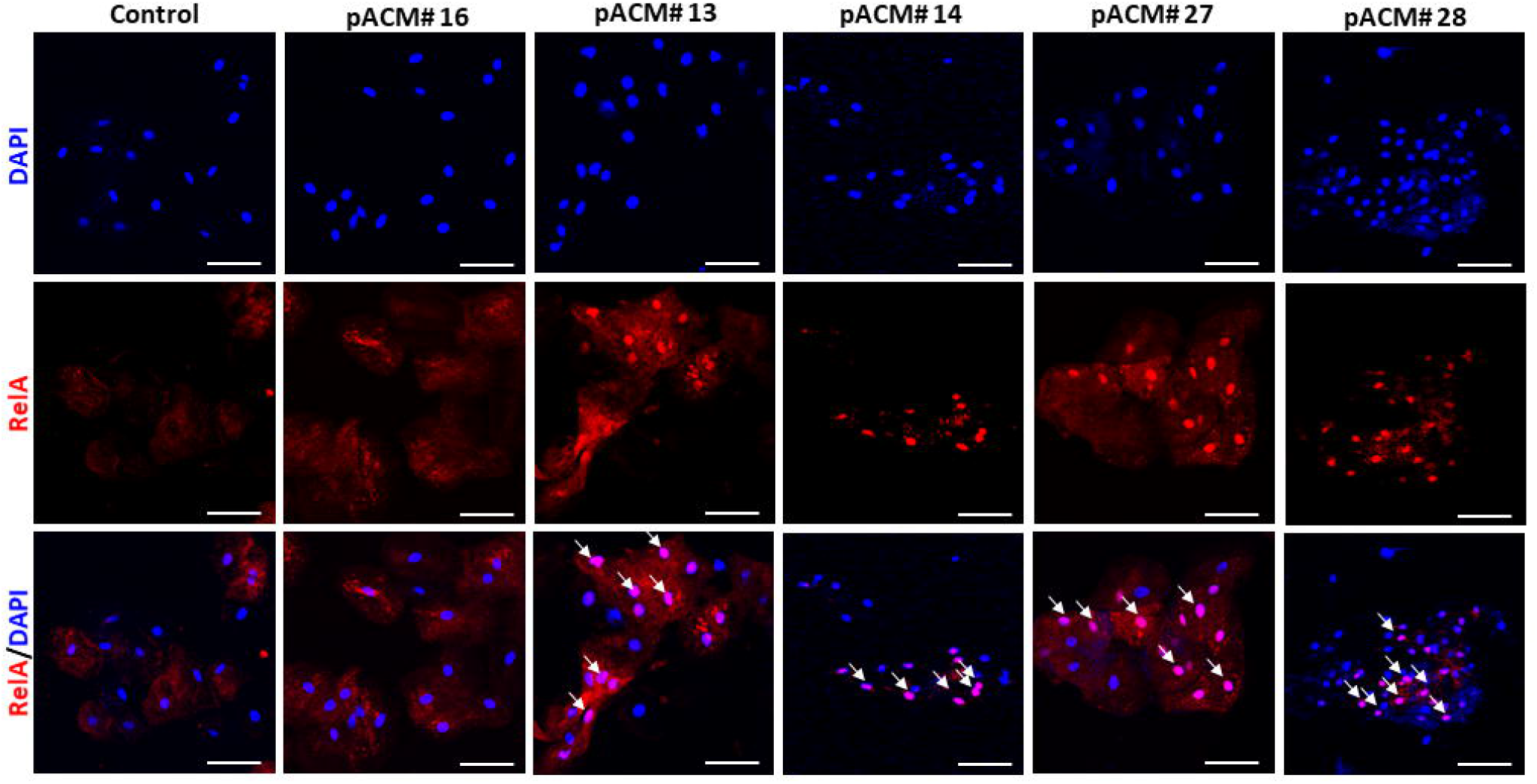
Expression of RelA/CCR2 in patient myocardial samples. **A**. Representative RelA/p65 immunoperoxidase staining in myocardial samples from controls and ACM patients. Brown nuclear signal (red arrows) in cardiac myocytes occurred only in ACM samples (x40); size bars, 50μm. **B**. Representative CCR2 immunofluorescent staining in myocardial samples from controls and ACM patients. White arrows, CCR2-immunoreactive signal was seen in small interstitial cells (x20); size bars, 100μm. **C**. The number of CCR2+ cells/mm^2^ section area in control and ACM patient heart samples; mean ± SEM; * p<0.0001 vs. controls; 1-way ANOVA with Tukey multiple comparison test. **D**. The number of CCR2+ cells vs. RelA+ myocyte nuclei/mm^2^ section area in n=11 ACM and n=12 controls; * p<0.0001 vs. controls determined by Pearson’s r correlation with 95% confidence intervals (dashed lines).

### CCR2+ cells are increased in hearts of ACM patients

In all control and ACM cases in which sufficient tissue was available, the number of CCR2+ cells per unit section area was determined by counting cells showing strong immunoreactive signal for CCR2 (**Figure 1B**). ACM hearts contained 15.2 ±4.1 CCR2+ cells/mm^2^ section area compared with 1.7 ± 1.4 for control hearts (p<0.0001; **Figure 1C**). Cells showing positive signal were small and round, and located between cardiac myocytes, consistent with their being macrophages (**Figure 1B**).

Our studies in *Dsg2*^*mut/mut*^ mice showed that NFκB signaling in cardiac myocytes was responsible for mobilizing CCR2+ cells to the heart where they promoted myocardial injury and arrhythmias (4). Accordingly, we compared the average number of cardiac myocyte nuclei showing immunoreactive signal for RelA/p65 and the average number of CCR2+ cells per mm^2^ section area in each case. As shown in **Figure 1D**, there was a close correlation, consistent with the causal mechanistic relationship observed in *Dsg2*^*mut/mut*^ mice (4).

### NFκB signaling is activated in buccal mucosa cells in young ACM patients with active disease

Buccal cells from the 12 ACM gene carriers with no clinical evidence of disease and the 9 patients with established but stable, quiescent disease showed no nuclear signal for RelA/p65, nor was nuclear signal seen in buccal cells from any of the 22 controls. By contrast, all 7 ACM gene carriers sampled either when clinical disease was first manifest or during “hot phases” showed strong RelA/p65 signal in buccal cell nuclei. Representative images are shown in **Figure 2**. Two subjects who showed nuclear RelA/p65 signal during a “hot phase” exhibited no nuclear signal in previous buccal smears analyzed 7 and 12 months before the onset of the “hot phase” (pACM27 and pACM26, respectively, in Table 2). Another patient with stable quiescent disease (pACM13) showed loss of nuclear signal in buccal cells analyzed 20 months after initial disease manifestation, whereas nuclear signal persisted in a repeat sample in a patient (pACM28) who had presented with a dramatic arrhythmia burden 2 months earlier, despite reduced arrhythmias after amiodarone therapy.

**Figure 2:**
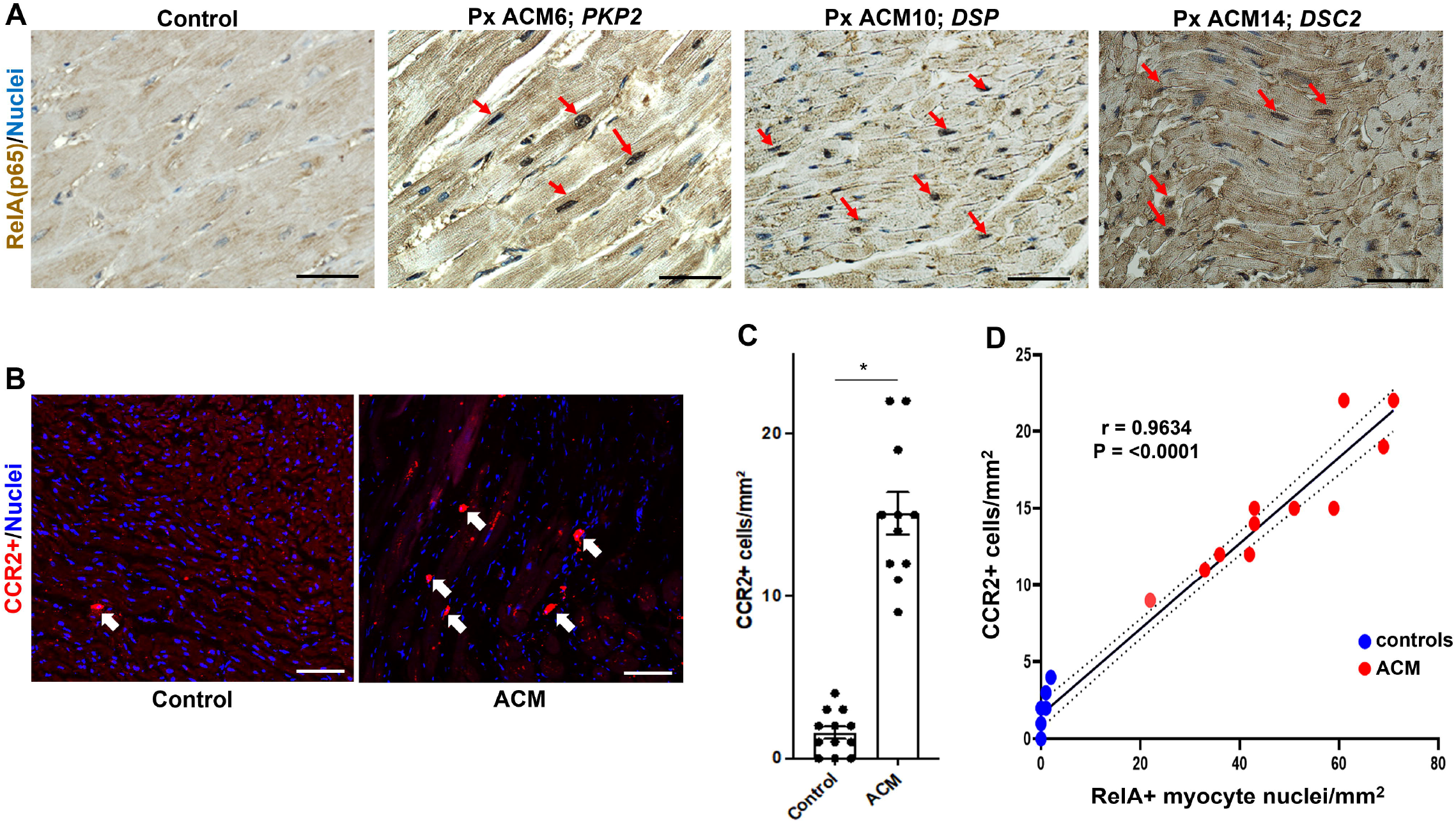
Expression of RelA in patient buccal smears. Representative RelA/p65 immunofluorescent staining in buccal cells from a control subject and pediatric ACM patients including those with stable disease (pACM# 16), at initial manifestation of disease (pACM# 13, pACM# 14) or during a “hot phase” of disease deterioration (pACM#27, pACM# 28). Nuclear RelA signal (white arrows) occurred only in patients undergoing phenoconversion or “hot phases”. Cell nuclei (blue) were counterstained with DAPI (x40); size bars, 50μm.

## DISCUSSION

We have previously reported that inhibition of NFκB signaling rescues the disease phenotype in *Dsg2*^*mut/mut*^ mice (3). NFκB signaling is also activated in iPSC-cardiac myocytes derived from patients with disease-causing variants in *PKP2* and *DSG2* (3, 8). These cells produce and secrete large amounts of pro-inflammatory mediators under basal conditions in vitro without prior stimulation or provocation. Taken together, these results suggest that innate immune signaling in ACM is a persistent, cell-autonomous inflammatory process that fails to resolve.

Here, we provide evidence that NFκB is activated in cardiac myocytes in ACM patients. We observed nuclear signal for RelA/p65 in cardiac myocytes, a reliable indicator of active NFκB signaling, in hearts of ACM patients who died suddenly and in others with advanced disease requiring cardiac transplantation. These observations suggest that as in *Dsg2*^*mut/mut*^ mice (4), ACM patients with clinically active disease exhibit a persistent innate immune response that does not resolve. We also provide new evidence linking NFκB signaling in cardiac myocytes with accumulation of CCR2+ cells in the hearts of ACM patients. These results suggest that persistent innate immune signaling in cardiac myocytes may be a major driver of disease in AMC patients, as occurs in *Dsg2*^*mut/mut*^ mice. If so, then ACM patients with clinically active disease might benefit from anti-inflammatory therapy that blocks NFκB signaling, as we have demonstrated in *Dsg2*^*mut/mut*^ mice (3). In addition, imaging methods to quantify myocardial CCR2+ cells might also identify patients at greatest risk of adverse events in ACM.

We also show here that NFκB signaling is activated in buccal mucosa cells of young ACM gene carriers at the time of disease onset or exacerbation. This observation extends previous findings indicating that ACM patients have a cutaneous phenotype, albeit subclinical, in which non-keratinized buccal epithelial cells exhibit features similar to those seen in cardiac myocytes in ACM (9, 10). It remains unexplained how ACM alleles persistently activate innate immune signaling pathways, but our findings suggest that inflammation in ACM extends beyond the heart, consistent with previous studies showing elevated circulating levels of pro-inflammatory cytokines in ACM patients (11). It is worth emphasizing that these studies would never have been possible by analyzing endocardial biopsies from young ACM gene carriers, whereas obtaining repeat buccal mucosa samples even from young children is safe, painless and well-tolerated by patients and their families. Using this approach, we observed that NFκB signaling is activated in buccal cells concomitant with onset of signs and symptoms ACM and persists in patients with clinically active disease. In view of the ease of obtaining and analyzing buccal smears, screening these cells for evidence of immune activation may help identify patients who would benefit from anti-inflammatory therapies.

## PERSPECTIVES

### Translational outlook

Patients with arrhythmogenic cardiomyopathy (ACM) develop progressive ventricular dysfunction and are at risk for sudden death. No mechanism-based therapies are available. Implantable defibrillators for high risk patients may be life-saving, but they do not treat the underlying disease or limit progressive myocardial injury. Compelling evidence from experimental models shows that persistent innate immune signaling via NFκB in cardiac myocytes causes arrhythmias and ongoing myocardial damage in ACM, at least in part, by mobilizing injurious macrophages expressing CCR2 to the heart. Here, we provide new evidence that NFκB signaling is activated in cardiac myocytes in hearts of ACM patients who died suddenly or who came to cardiac transplantation. This was associated with a marked increase in the number of infiltrating CCR2-expressing cells in their hearts. We also show that active NFκB signaling occurs in buccal mucosa cells of young disease allele carriers at the time of their initial onset of disease or after experiencing disease exacerbation. These results suggest that anti-inflammatory drugs to block persistent NFκB signaling may be an effective mechanism-based treatment for ACM patients. Screening buccal cells for NFκB signaling may be a safe, inexpensive and reliable way to prospectively identify ACM patients who might benefit from anti-inflammatory therapy.

## ABBREVIATIONS

ACM: arrhythmogenic cardiomyopathy
pACM: pediatric arrhythmogenic cardiomyopathy
NFκB: nuclear factor κB
CCR2: C-C motif chemokine receptor-2
MCP1: monocyte chemoattractant protein-1
iPSC: induced pluripotent stem cells
DAB: 3,3’-diaminobenzidine
DAPI: 4’,6-diamidino-2-phenylindole
SCD: sudden cardiac death
PM Dx: post-mortem diagnosis
RV-dom: right ventricular dominant
LV-dom: left ventricular dominant
HF: heart failure
ECG: electrocardiogram
ICD: implantable cardiac defibrillator
cMRI: cardiac magnetic resonance imaging
LGE: late gadolinium enhancement
NSVT: non-sustained ventricular tachycardia
SVT: sustained ventricular tachycardia
PVC: premature ventricular complex
SAECG: signal-averaged electrocardiogram

## ACKNOWLEDGMENTS

We thank the patients and their families who participated in this study.

